# Adhesion modulates cell morphology and migration within dense fibrous networks

**DOI:** 10.1101/838995

**Authors:** Maurício Moreira-Soares, Susana P. Cunha, José Rafael Bordin, Rui D. M. Travasso

## Abstract

One of the most fundamental abilities required for the sustainability of complex life forms is active cell migration, since it is essential in diverse processes from morphogenesis to leukocyte chemotaxis in immune response. The movement of a cell is the result of intricate mechanisms, that involve the coordination between mechanical forces, biochemical regulatory pathways and environmental cues. In particular, epithelial cancer cells have to employ mechanical strategies in order to migrate through the tissue’s basement membrane and infiltrate the bloodstream during the invasion stage of metastasis. In this work we explore how mechanical interactions such as spatial restriction and adhesion affect migration of a self-propelled droplet in dense fibrous media. We have performed a systematic analysis using a phase-field model and we propose a novel approach to simulate cell migration with Dissipative Particle Dynamics (DPD) modelling. With this purpose we have measured the cell’s velocity and quantified its morphology as a function of the fibre density and of its adhesiveness to the matrix fibres. Furthermore, we have compared our results to a previous *in vitro* migration assay of fibrosacorma cells in fibrous matrices. The results are model independent and show good agreement between the two methodologies and experiments in the literature, which indicates that these minimalist descriptions are able to capture the main features of the system. Our results indicate that adhesiveness is critical for cell migration, by modulating cell morphology in crowded environments and by enhancing cell velocity. In addition, our analysis suggests that matrix metalloproteinases (MMPs) play an important role as adhesiveness modulators. We propose that new assays should be carried out to address the role of adhesion and the effect of different MMPs in cell migration under confined conditions.

## 1. Introduction

Cancer progression is a multistep process. During metastasis in solid carcinomas, tumor cells leave the primary colony, invade the circulatory system, reach distant organs and form new cancer cell colonies. This movement of cancerous cells to other organs and tissues is the main cause of death in oncological patients [1, 2]. The existence of metastasis is one of the markers that indicate poor prognosis [3, 4]. This intricate mechanism can often be divided into distinct stages: invasion, intravasation, extravasation and colonisation [1, 5]. In this work we focus on the invasion stage, described by the migratory event that takes place after the detachment of a tumor cell from the primary colony. As the tumor cells leave the primary tumor and start their journey in the direction of a blood vessel there are signalling cascades and mechanical regulation events that modulate cellular migration [6, 7, 8]. In the last decades, due to the advance of microscopy and computational modelling, experimentalists and computational biologists have been studying the mechanical aspects of cellular behaviour with the aim of better understanding the complex process that is metastasis [9, 10, 11, 12, 13, 14]. However, the current state of the art is still far from the full comprehension of the mechanics behind cancer cell migration in tissues *in vivo*.

One of the main regulators of cell function is the microenvironment, and in particular the Extracellular Matrix (ECM). The ECM is a rich network of macromolecules that occupies the interstitial space between cells and provides biochemical and mechanical support [15]. The ECM is composed of water, proteoglycans and fibrous proteins such as collagen, as well as signalling molecules and growth factors [6, 16]. This structure can be remodelled and degraded by cells through their mechanical action or by chemical reactions [17, 18]. Enzymes present in this environment, among other functions, regulate and/or promote the proteolytic activity within the ECM while reshaping the medium [19, 20]. The biochemical content and the mechanical properties of the ECM may vary from tissue to tissue, affecting the behaviour of the cells that are dependent of these properties [16]. In particular, the mechanical constraints imposed by the ECM may affect directly cell migration [21, 22]. For instance, high density collagen matrices induce the cells to employ new strategies in order to enhance migration within the restricted space environment [23, 24, 25]. There are several experimental observations which suggest that cells in confined space migrate preferentially along fibronectin fibres and preexisting paths in the ECM [26, 27], searching for the direction of least resistance [28, 29, 30, 31] and deforming their shape in order to follow these channel-like structures [32, 33].

The cells’ capability of adhering to their surrounding is mediated by transmembrane proteins in the cell membrane. These matrix receptors bind proteins present in the ECM to form cell-matrix junctions [34]. The main group of cell adhesion molecules (CAMs) responsible for linking the cell to the matrix components is the integrin family. Integrins are able to form anchoring junctions linking actin filaments and intermediate filaments in the cell interior to the collagen, fibronectin and laminin from the ECM [35]. To increase adhesion it is then required the clustering of integrins with the purpose of forming the anchorage of the different filaments.

There are different kinds of adhesion structures with specific functions: focal adhesions (FAs), responsible for strong adherent sites during cell migration; fibrillar adhesions, that are originated from the continuous application of forces through FAs, depending on actomyosin contractility and matrix bending rigidity [36]; podosomes/invadopodia, which are actin rich structures and distinguishable from FAs due to their shorter lifespan of a few minutes and their additional function in promoting invasion in the ECM [37, 38], among others.

The matrix metalloproteinases (MMPs) belong to a family of enzymes that catalyse the cleavage of proteins [39] and initially emerged as the main collagen-degrading enzymes. Since then, further studies have demonstrated their proteolytic influence over other components of the ECM [40, 41]. For a long time the importance of the MMP family was restricted to this single role and consequently obscured its crucial position in cell dynamics. In fact, MMPs act also by regulating cell functions in a broader sense through the cleavage of others proteins, which have consequences in cell adhesion, proliferation, survival and migration. MMP activity is linked to the cell ability of adhering to the ECM due to its influence on specific CAMs [42, 43, 44], and even to the regulation of actin polymerisation [45].

Mathematical modelling has demonstrated its usefulness in the study of biological systems when coupled with experimental assays [46, 47]. In our perspective, the aim of building a mathematical description for such intricate systems is not to completely characterise the system in its full complexity. Instead, the challenge is to implement assertive assumptions and approximations that simplify the study while preserving the key features of the biological system. Strikingly, recent works have adopted two or more different mathematical approaches to tackle the same problem in order to test whether the common mechanistic assumptions of each model are relevant or not [48, 49]. When different models of a single biological process yield similar behaviours then the results are robust to the modelling implementation. In this work we employ two particular approaches for cell migration that are complementary to each other, the phase-field model (PFM) and the dissipative particle dynamics (DPD).

Phase-field models have been extensively used to study biological systems, such as in tumor growth [50, 51, 52, 53], vessel growth [54, 55, 56, 57], cell monolayers [58, 59], axonal development in neurons [60], immune system response [61, 62] and cellular motility [63, 64, 65, 66, 67, 68]. Since they are focused on the interface dynamics, a major advantage of PFMs is the lower number of parameters when compared to other established methods such as Cellular Potts Models (CPMs), Agent-Based Models (ABMs) or mixture models. As a result, this low number of parameters gives the possibility to build minimalist models to approach specific questions about the system of interest. In particular, in this work we model a self propelled cell as a droplet within a porous material composed of a fibre network. The cell maintains its shape due to the surface tension, and interacts with the fibres by adhesion and depletion. We carry out a systematic study by varying two parameters: the density of the fibres and the adhesion strength between the cell and the fibres.

Recently, the DPD approach has been used in the computational biology field to describe tissues and cells [69, 70, 71, 72]. The DPD model is a particle-based method introduced in 1992 for hydrodynamics phenomena simulation which combines the algorithmic scheme of Molecular Dynamics (MD) and the time-stepping of Lattice-Gas Automata (LGA) [73]. These features delivered a method faster than MD and more flexible than LGA, thus enabling the exploration of systems with larger spatial scale and for longer timescales. Similar to the aforementioned methods, DPD uses Newtonian mechanics for calculating explicitly the interaction forces between particles. In this work we introduce a novel description for modelling the cell membrane and the ECM using DPD. Similarly to the phase-field model, our system is composed by the cell and the fibres. The cell is described as a spherical distribution of monomers around a central monomer at a fixed distance. The monomers at the cell surface are linked to the central monomer through a harmonic potential, mimicking an elastic membrane. On the other hand, the fibres are modelled as chains of monomers connected via a harmonic potential and randomly distributed in space.

In the following section, we present the two computational methods adopted and the mathematical details behind each approach. In the section Results and Discussion, we compare the results between the two models. We also compare our results with experimental data available in the literature and, finally in the Conclusions section, we draw some conclusions and introduce hypotheses about the biological processes modelled with these systems.

## 2. Methods

### 2.1. Phase-Field Model

Our simulation is based on the mathematical model for multicellular systems introduced in ref. [74] and explored further in [75]. First we define a free energy functional *F* [*ϕ, ψ*] that will determine the system dynamics. Similarly to [56], we use two order parameters *ϕ*(**r**, *t*) and *ψ*(**r**, *t*). The cell is identified in space by the regions where *ϕ* ≈ 1 and the ECM is defined by the domain where *ψ* ≈ 1 (see Fig.1). The interstitial space is defined as the region where both functions are close to zero. This free energy functional is the sum of a functional related to the cell’s energy *F*_*cell*_[*ϕ*], with another functional describing the cell’s interaction with other biological entities *F*_*int*_[*ϕ, ψ*], such as the ECM:.

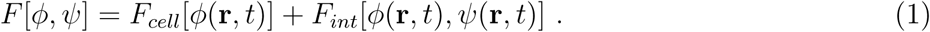

The first component of this functional, written in Eq. 2, is given by the Ginzburg-Landau free energy with a double well potential, with two stable solutions *ϕ* = {0, 1}, which correspond respectively to the exterior and interior of the cell (see Fig. 2),

**Figure 1.**
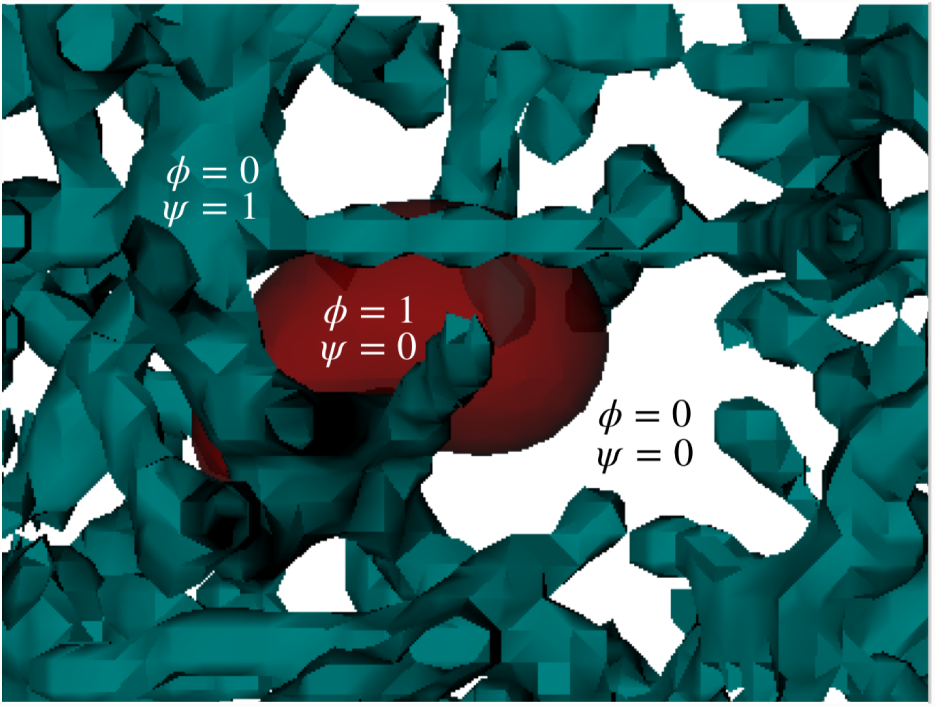
The fields *ϕ* and *ψ* are depicted in the image. The cell (red) is defined by the region where *ϕ* = 1 and *ψ* = 0, while the fibres (cyan) are represented by the domains of *ψ* = 1 and *ϕ* = 0. The interstitial space is represented by both fields being zero simultaneously (white).

**Figure 2.**
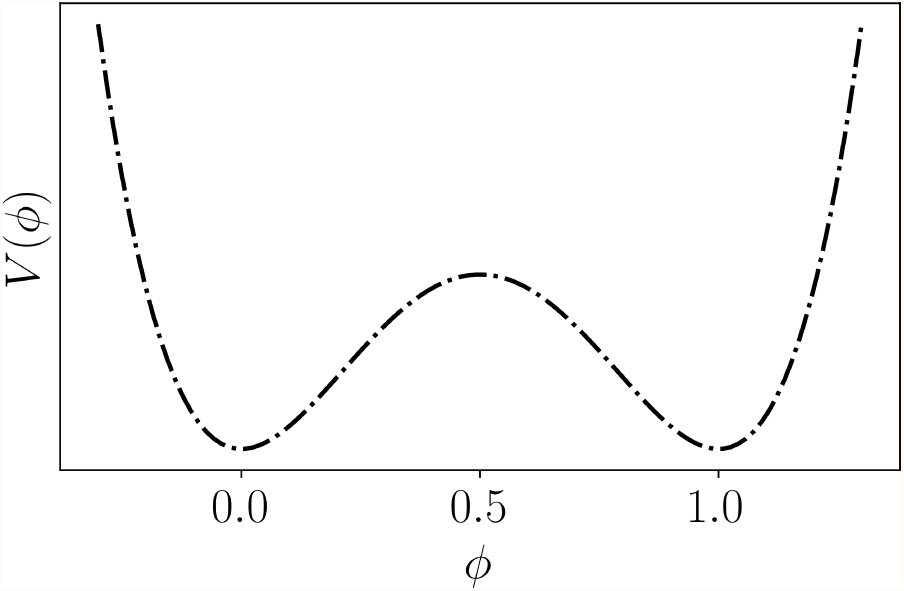
The double-well potential is used in order to maintain the two stable phases for describing the cell interior and the exterior. The two minima are symmetrical and at *ϕ* = 1 and *ϕ* = 0 respectively [76].

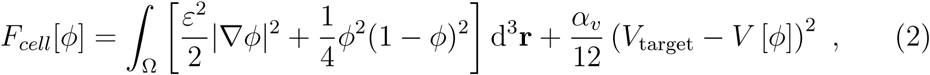

where Ω denotes the total volume of the system. The coefficient *ε* is a positive constant that is related to the width of the interface and to the surface tension *σ*. The squared gradient term acts as an energetic cost for maintaining the interface: more interface costs more energy. The last term is the Lagrange multiplier for the volume *V* [*ϕ*] = ∫ _Ω_ *h*(*ϕ*)d^3^ **r**, where *h*(*ϕ*) = *ϕ*^2^(3 − 2*ϕ*). The function *h*(*ϕ*) reinforces the order parameter values to remain between the interval [0, 1], since *h*′(*ϕ*) = 0 for *ϕ* = 0 and *ϕ* = 1.

The interaction component of the free energy is given by

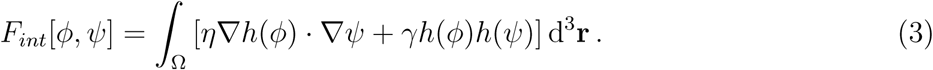

The first term characterises the adhesion between the cell and the matrix (controlled by the parameter *η*), and the second penalises energetically the overlap between cells and matrix fibres (controlled by the parameter *γ*). Given the free energy functional, the equation for the cell dynamics is derived as a gradient descent equation with an advective term. Therefore, in this case we consider the *ϕ* change rate to be proportional to the gradient of the free energy functional [76, 77]

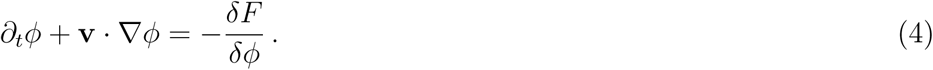

Taking the free-energy functional derivative, we finally have the equation for the cell dynamics

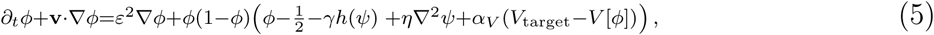

where the velocity **v** is written as the gradient of the chemical field *c*, which for metastatic cells can be assumed as the oxygen concentration in the tissue guiding the cell to a blood vessel during invasion:

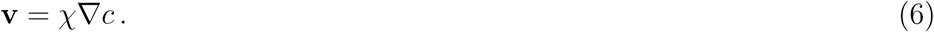

In this study we assumed that this velocity is constant and directed along the *x*–direction, which allowed us to isolate the influence of the mechanical constraints in the movement. Furthermore, we also tested introducing a white noise in the velocity using a Persistent Random Walk (PRW) model and the results remained robust to this alteration. The details regarding the numerical implementation are in the Supporting Information (SI) and the software is available in the online repository https://phydev.github.io/SPiCCAto [78, 79].

### 2.2. Dissipative Particle Dynamics

We propose a model for the system described in the previous sections using a DPD approach. Moreover, the DPD allows us to explore different perspectives that would be extremely complicated using only the PFM, such as the introduction of elastic fibres. Briefly stated, in DPD we solve Newton’s equation of motion based on the total force that acts on each particle *i* [70, 80]:

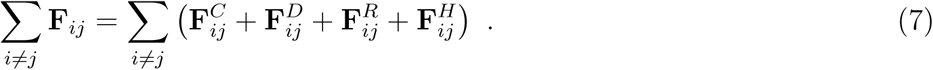

The first term in this equation is the conservative force, which models the interaction between the different bead types, as function of the beads’ types. Here, our conservative term is

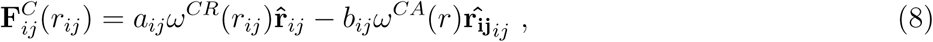

where the first term is the repulsive force, with *a*_*ij*_ being the maximum repulsion between the particles *i* and *j*, and the second term the attractive force with maximum intensity *b*_*ij*_. In Eq. 8 **r**_*ij*_ = **r**_*i*_ − **r**_*j*_ is the distance between the two particles, and *ω*^*CR*^(*r*), known as the weight function, usually assu es the form

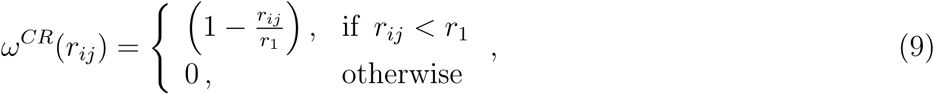

with *r*_1_ = 1.25 *µ*m. This term is responsible for the repulsion due the excluded volume of the beads. To model the adhesion of cells to the ECM fibre we added the second term of Eq. 8. This attractive term in the conservative force was proposed recently for the study of cell deformation in a surface [81]. The attractive weight function is given by:

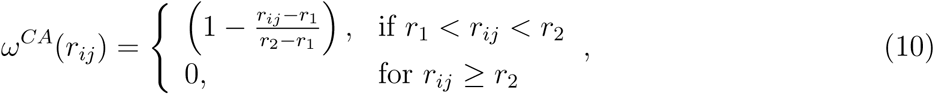

where *r*_2_ = 2.50*µ*m. This term acts only near the fibres’ surface and describes the biochemical adhesion forces between the cell and the fibres.

The dissipative force, 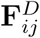 in Eq. 7 represents the effect of viscosity. It uses the relative velocity between two particles to slow down the motion with respect to each other and is given by

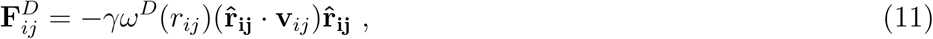

where *γ* is a coefficient, **v**_*ij*_ = **v**_*i*_ − **v**_*j*_ is the relative velocity between the particles and *ω*^*D*^(*r*_*ij*_) is a weight function that depends on the distance between the particles:

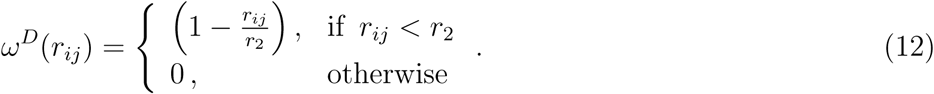

Finally, the random force 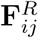 represents the thermal or vibrational energy of the system and includes fluctuation effects,

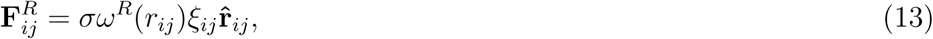

where *σ* is a coefficient, *ω*^*R*^(*r*_*ij*_) the weight function and *ξ*_*ij*_ a random constant obtained from a Gaussian distribution.

The dissipative and random forces are related by their coefficients and weight functions,

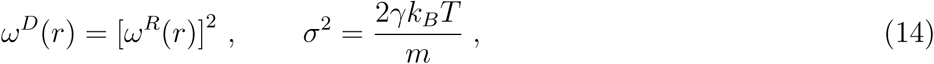

where *k*_*B*_ is the Boltzmann constant and *T* the equilibrium temperature. These relations turn the DPD into a thermostat. Since the algorithm depends on the relative velocities between the beads and since all interactions are spherically symmetric, the DPD is an isotropic Galilean invariant thermostat which preserves the hydrodynamics [82].

Here, we propose a model to study the relation between cell shape and dynamics. Our cell consists of 452 beads distributed around a sphere of radius *a*_*c*_ = 6*µ*m, the cell radius. Cell surface beads *c* repel themselves by the conservative force with an intensity of *a*_*cc*_ = 12.5 × 10^−11^ N. These surface cell beads are attached by a standard harmonic force **F**^*H*^ to a central ghost bead *g*, as depicted in Fig. 3(a):

**Figure 3.**
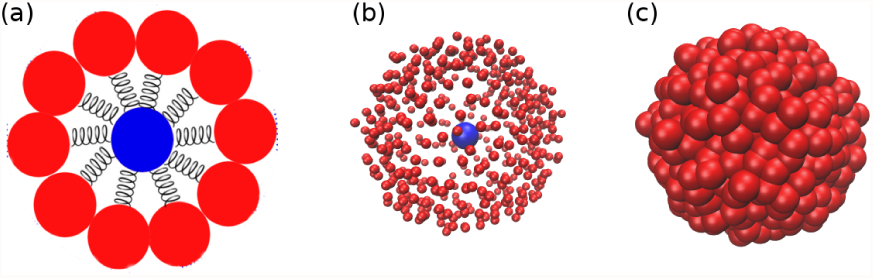
(a) Schematic depiction of the DPD cell model. (b) Cell in a bulk simulation showing the central ghost bead. (c) Same as (b), but with the surface beads with the correct size that is included in the simulation.

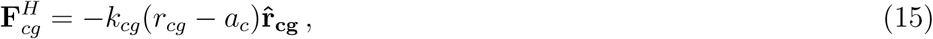

where **r**_**cg**_ is the distance between a *c* bead and the central bead *g*, and *k*_*cg*_ = 30 N/*µ*m. Fig. 3(b) shows a snapshot of the cell in a bulk simulation with small surface beads so the ghost bead can be seen. In Fig. 3(c) we show the same snapshot but with the surface beads depicted in their regular size. As we can see, the cell remains approximately spherical - obviously less spherical than the raspberry model [83] - and, as we will demonstrate, it can deform to pass through obstacles.

The other two species in the DPD system are the solvent and the fibre matrix. Solvent beads *s* have a number density *ρ*_*N*_ = 4.0, consistent with previous works [84, 85], and are repelled by *c* beads with a strength *a*_*cs*_ = 65 × 10^−11^ N with a cut off radius of 2.5 *µ*m. The fibres are modelled as polymers with beads of type *f*, and the interaction parameters between *c* and *f* beads are the same as between *c* and *s, a*_*cs*_ = *a*_*cf*_. The only attraction interaction in the system is between *c* and *f* beads. In this case, the attractive parameter plays the role of the adhesion defined by the relation *b*_*cf*_ = *η*_*DPD*_*a*_*cf*_. Here *η*_*DPD*_ provides the strength of the adhesion between the cell and the polymer network.

In the case of a rigid fibre network, the polymers are placed randomly inside the simulation box and remain fixed, i.e., the corresponding beads are not integrated in time. In the case of flexible fibre network only the beads placed at the ends of the polymeric chains are fixed, and the rest of the polymer can oscillate similarly to a rope with fixed ends. In this flexible case, the beads in the same polymer are bound by the harmonic bond, Eq. 15, with *k*_*ff*_ = *k*_*cg*_ = 30 N/*µ*m. The case of rigid polymers can be imagined as the limit case of *k*_*ff*_ → ∞.

The cell is placed in the left buffer zone of the simulation box and a constant force in the longitudinal direction is applied to the cell mimicking the chemical gradient. Then we analyse the system properties during the cell migration in the fibre network region. The simulation ends when the cell reaches the right buffer. The SI presents a detailed description of the setup. We show in Fig. 4 a schematic depiction of the simulation box.

**Figure 4.**
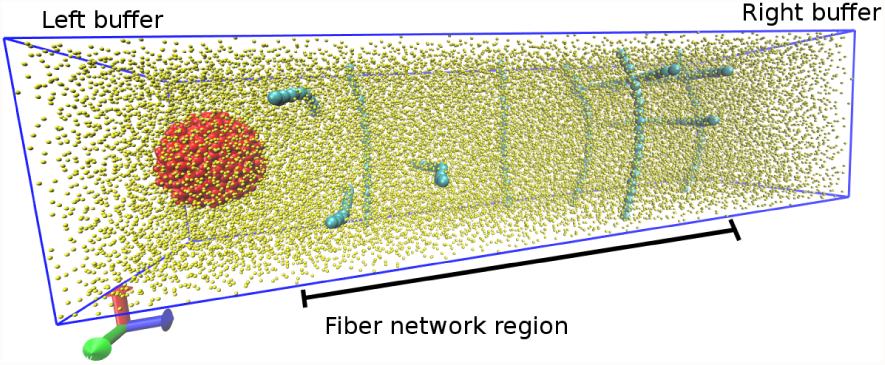
Depiction of the DPD model for a flexible network. The fibres (cyan) are represented by a chain of monomers connected by a harmonic potential. The cell (red) is modelled as an approximately spherical set of beads bound by an elastic potential to a central bead inside the cell. The solvent is represented in yellow. For better visualisation the solvent beads are smaller than the other beads and only a few fibre chains are shown.

## 3. Results and Discussion

As previously described in Section 2, we start by simulating a droplet migrating trough the fibre network using both the PFM and DPD model, and quantify the influence of the medium in its shape and movement. As expected, the properties of the fibre network, particularly the pore cross section, adhesiveness and elasticity, affect cell migration and morphology.

We quantify migration by measuring the mean velocity *(v)*, the mean square displacement (MSD) and the diffusion exponent *β*. In addition, morphology is quantified by the cell’s radius of gyration *R*_*g*_ and surface energy, which measure deformation and the surface roughness respectively. These properties are obtained as a function of the fibre density *ρ*, defined by the quotient of occupied and total volume of the system, the mean pore cross section (see SI) and the adhesion strength *η* between the droplets and the fibres. We compare our results to experimental data for cell migration *in vitro*.

### 3.1. Cell movement and matrix density

For the cell to overcome the obstacles in dense fibre networks it needs to deform. In Fig. 5 we show some snapshots from the cell migration. Row (A) refers to cell modelled with DPD when the fibres are rigid, while row (B) shows examples for the flexible matrix scenario. For better visualisation, the fibre beads are smaller than the cell beads, and the solvent beads are not shown. Comparing (A.I) and (B.I) snapshots, we can see that even for a simple obstacle, which is to bypass a single fibre, the cell is more deformed by the rigid fibre. The deformation becomes larger when the cell is blocked by more than one rigid fibre, as the (A.II) and (A.III) snapshots show. On the other hand, the cell is only slightly deformed by the flexible matrix, as the (B.II) and (B.III) snapshots show. We expect that the time needed by the cell to escape from a fibre trap is directly related with the time necessary to deform its spherical shape and cross through the available space. The PFM presents similar results to the DPD with rigid fibres, which we can see when comparing the rows (A) and (C) in Fig. 5.

**Figure 5.**
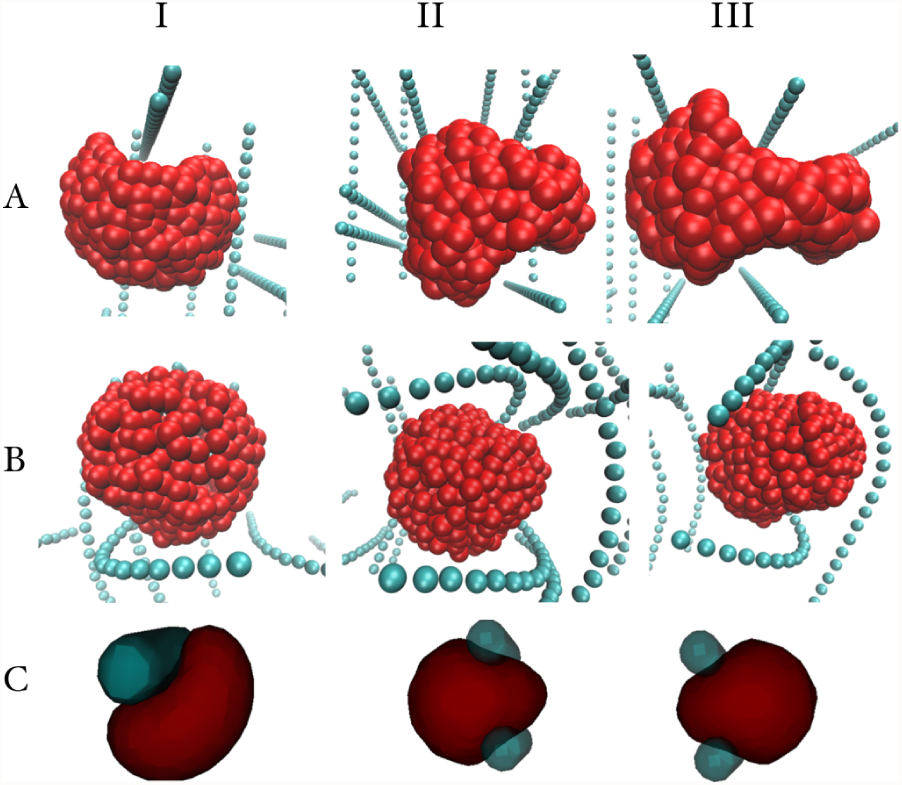
Examples of cell deformations inside rigid (**A**) and flexible (**B**) fibres in DPD compared to phase-field model (**C**).

To understand the velocity dependence on the fibre density *ρ* for rigid and flexible networks we present in Fig. 6 typical examples of the *x* component of the cell center of mass (CM) of the DPD cell as function of time for three different densities *ρ* = {0.20, 0.55, 0.70}. We present similar results for the PFM in Fig. 3 of the Supporting Material.

**Figure 6.**
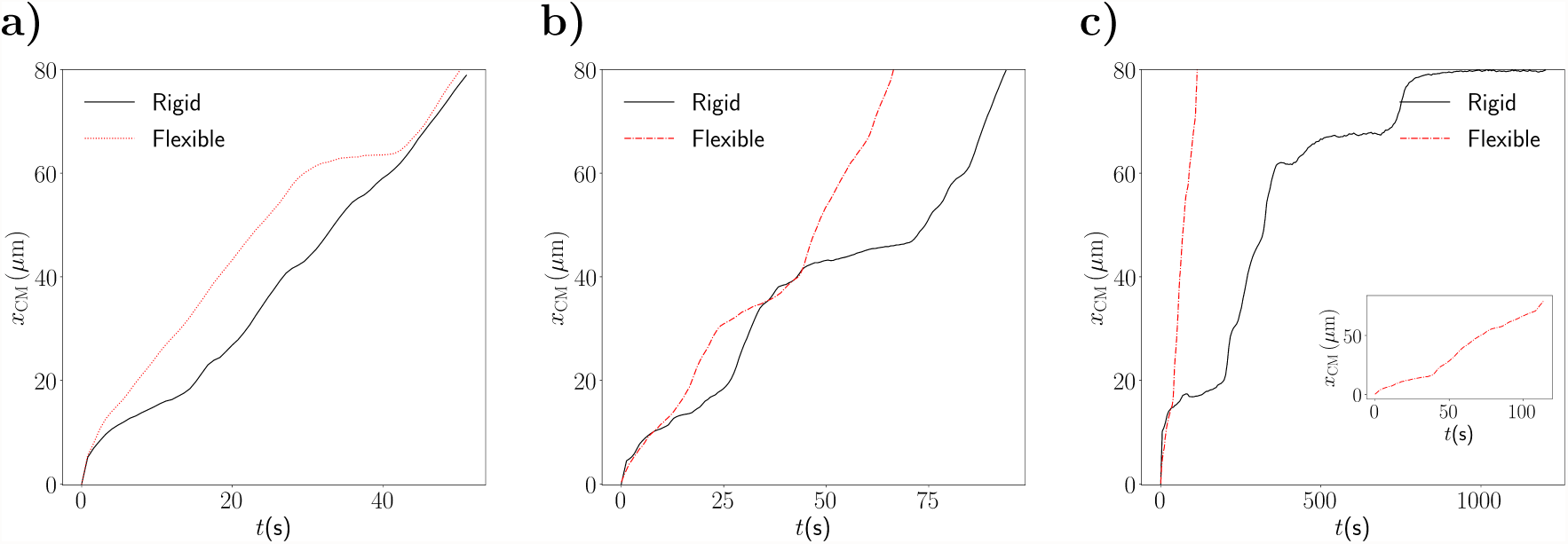
Position of the cell center of mass as function of time during its movement for fibre densities **a)** *ρ* = 0.20, **b)** *ρ* = 0.55 and **c)** *ρ* = 0.70. The inset in **c)** shows the curve for the flexible matrix.

In the examples for the lowest fibre density, the cell may be blocked a few times for both rigid and flexible fibre matrices, but it diffuses almost without obstacles for most of the time. In these low density matrices, the time required for the cell to leave the rigid or flexible network is similar. However, as *ρ* increases the scenario changes. For *ρ* = 0.55, Fig. 6(b), we can see that both curves have two trap regions, but the time spent by the cell to leave those regions is much longer in the rigid network. The difference in the stop time for the rigid and flexible matrices becomes even more pronounced for *ρ* = 0.70, Fig. 6(c).

Another interesting feature in these systems is the diffusion regime. We quantify the diffusion by scaling the MSD and the time, namely *< r*(0)*r*(*t*) *>*≈ *t*^*β*^. If the MSD increases linearly with time, *β* = 1.0, the system is in the normal or Fickian regime. If *β <* 1.0 the diffusion is anomalous and called subdiffusive, while in the case of *β >* 1.0 the regime is superdiffusive. The limit of *β* = 2.0 is called ballistic. To understand how the rigid or flexible matrix can influence the diffusion regime we evaluated the exponent *β* as function of the fibre density *ρ*. The value of *β* used here is the obtained as *t* → *∞*, and not the short time exponent.

We can observe in Fig. 7 that a superdiffusive regime was observed at lower values of *ρ* for the three models, in agreement with experimental studies [86]. However, at higher network densities, *ρ ≥* 0.50, we observe a change in the regime for the DPD model with rigid fibres and for the PFM. There is a transition from a superdiffusive to a subdiffusive regime. This transition can be directly related to the longer times the cell gets blocked the fibre matrix. At low densities, since there is force pulling the cell, the movement is almost ballistic, with occasional collisions with the matrix. However, as the density increases, the cell MSD is strongly affected, with long periods of no mobility. On the other hand, the migration inside flexible matrix is only slightly affected. The movement changes from a near-ballistic to a superdiffusive regime. Here, the change is small and even at high densities the time that the cell remains blocked is small, as we have showed in Fig. 6(c). Therefore, the flexibility of the fibre network plays an important role not only in the velocity of migration, but also in cell deformation and in diffusion regime.

**Figure 7.**
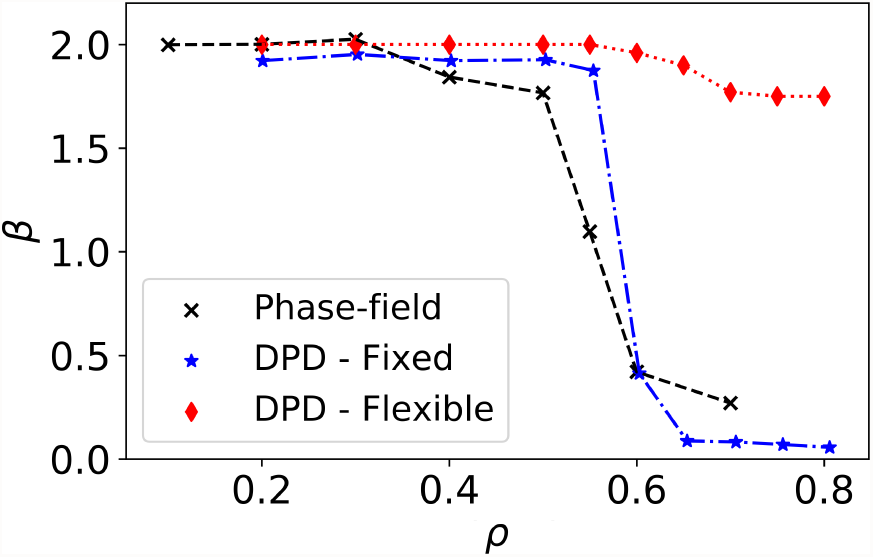
Exponent *β* as function of the fibre density *ρ*. For lower density of fibres (0.1 ≤ *ρ* ≤ 0.5) the exponent *β* remains unaltered with a value near 2, which indicates a ballistic behaviour present in the three models. When the density is increased even further a transition region appears for the PFM and for the DPD with fixed fibres, but not for the DPD with flexible fibres.

### 3.2. Adhesiveness as a key modulator in migration

We then explored how adhesion affects cell migration in both PFM and DPD models. The mean velocity is measured as the quotient of the trajectory traveled by the center of mass of the cell and its respective time interval. In Fig. 8 we present how the fibre density *ρ* affects migration for three different values of adhesion *η*. The overall behaviour is similar and independent of the adhesion coefficient: for lower fibre densities we obtain the highest velocities, while as the density increases, the velocity becomes smaller the cell gets trapped in the matrix. Nonetheless, by comparing the migration velocity for different values of adhesion at the higher densities *ρ* = {0.6, 0.7}, we observe that the velocity is higher for *η* = 1 than for the lower values *η* = {0.0, 0.5}. This dynamic is anticipated by experimental results of cell migration, but it is striking to observe this behaviour in this simple model.

**Figure 8.**
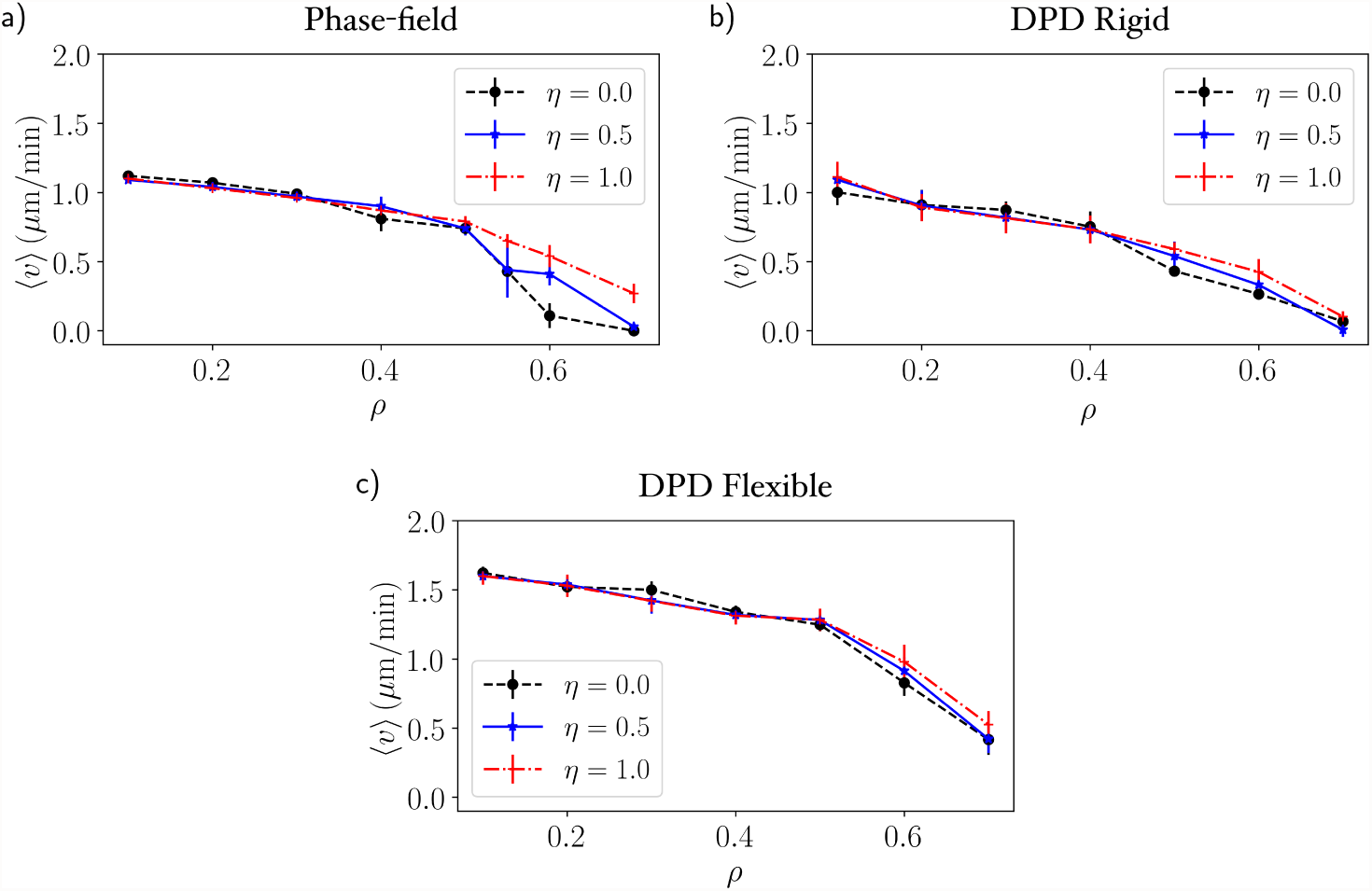
Migration velocity as a function of the density of fibres with three values of adhesion coefficient *η* = 0, 0.5, 1.0 for a) PFM, b) DPD with rigid fibres and c) DPD with flexible fibres. The velocity starts to decrease for *ρ >* 0.4 and the cells stop moving (except for the flexible fibres) at *ρ*_*M*_ ≈ 0.7.

For further investigation concerning the dependence of the migration velocity on the adhesion we fixed the density of fibres at *ρ* = 0.6 and ran simulations for several values of adhesion. In Fig. 9 the mean velocity is plotted as a function of the adhesion coefficient for our computational models and it includes the *in vitro* results adapted from [87]. We can see that the cell migration velocity increases with the adhesion coefficient until it reaches a maximum, after which it decreases. This suggests that there is an optimal value of adhesion for migratory events [88, 89]. Indeed, we have superposed the experimental data for the cell velocity as a function of the fibronectin concentration obtained by Maheshwari and colleagues [87] and the same qualitative behaviour is verified (see Fig. 9). Fibronectin appears as protagonist in cell adhesion and its concentration is a proxy for altering adhesiveness – more fibronectin present in the ECM leads to higher adhesiveness.

**Figure 9.**
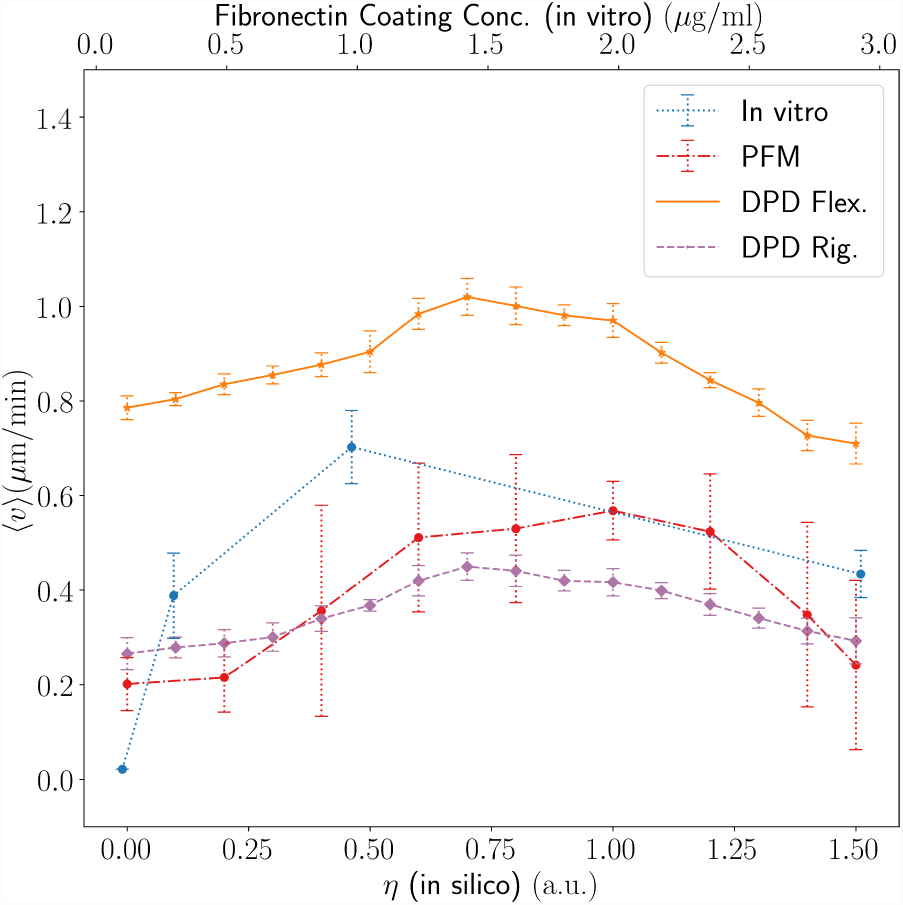
Migration velocity as a function of the adhesion coefficient for *in silico* results and as a function of the fibronectin coating concentration for the *in vitro* assay [87]. The fibronectin concentration is directly related to the adhesiveness, higher concentrations of fibronectin strengthen the adhesion between the cell and the ECM. The simulation results show clearly a peak in velocity around *η* = 1.0.

Experimentally the adhesiveness is linked to the cell capability to exert traction forces in the fibres in order to pull its body forward. Thus this is the main reason by which the *in vitro* assay exhibits an increase in velocity with the adhesiveness and then decreases after reaching a critical point where adhesion is so strong that it suppresses migration. However, in our mathematical model we have a slightly different circumstance that leads to the same qualitative results. The first main difference we should stress here is that adhesion is a passive attribute while in nature adhesion is an active dynamical event performed by the cell. The migration of our droplet is affected by adhesion in two different manners: i) changing cell plasticity and ii) introducing an artificial pulling force. A higher adhesion increases the cell deformity (see Fig. 10) thus facilitating the migration through small pores. Moreover, the adhesion acts as a pulling force due to the leading edge of the cell that feels the fibres upfront and is attracted to them by the adhesive force. This effect mimics some of the features observed in real cells during migration.

**Figure 10.**
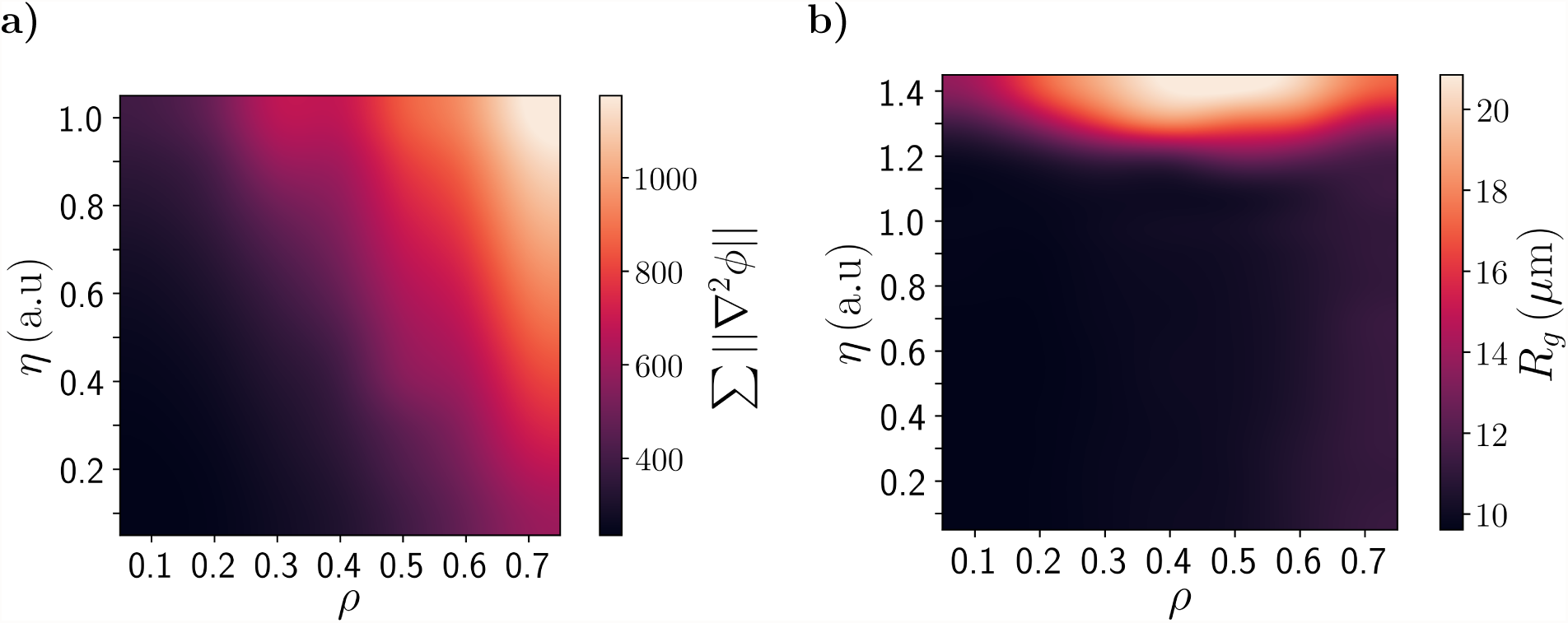
Surface energy (left) and radius of gyration (right) as function of the density of fibres and the adhesion coefficient. The surface energy gives an estimative of surface roughness and deformation. The surface energy increases with the adhesion with fibre density, which indicates a higher cell plasticity.

The study realised by Wolf K. et al. used an *in vitro* model of tumor cells and immune cells (T-cells and neutrophils) migrating in 3D collagen scaffolds of different porosities to study the physical limits of migration [9]. In order to compare our results to their experimental data, calculations were performed to determinate the mean pore cross-section of the fibre networks. Each matrix was sliced in several planes orthogonal to the migration direction and for each plane the Connected Component Algorithm (CCA) was applied to obtain the area of the pores (see Supporting Information). Then the procedure was repeated for several random generated matrices of the same density and the mean values and their respective unbiased standard deviation were obtained. On the other hand, for the DPD matrices we have used a method described in [90] that estimates the area of the pores by the effective space available [91]. The pore sizes is inversely proportional to the density of fibres. The computational data is depicted in Fig. 11 together with experimental results adapted from [9] (Fig. 2-B). The experimental data was measured for human fibrosarcoma cells of the line HT1080 migrating in rat tail collagen matrices treated with a MMP inhibitor (GM6001) and for untreated matrices (with medium).

**Figure 11.**
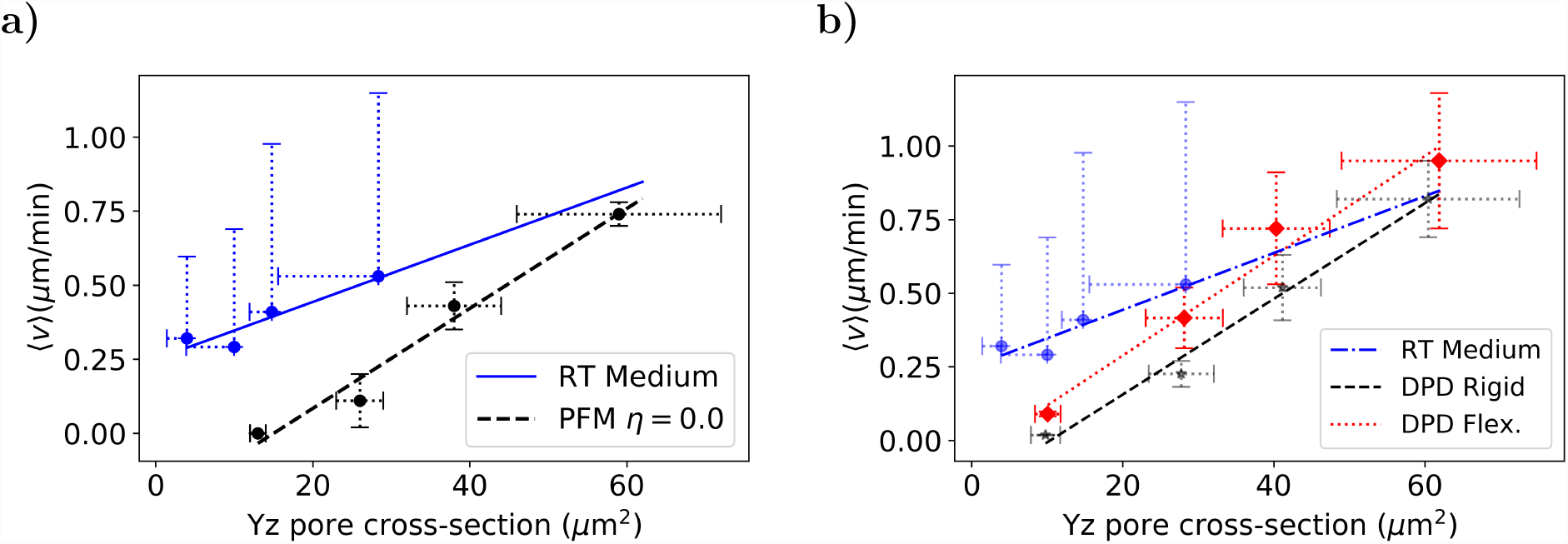
Migration velocity as a function of the pore cross-section *in silico* and *in vitro* for untreated rat-tail collagen [9]. The results for **a)** the phase-field model and **b)** the DPD model are similar. The slopes and *R*^2^ are the following: 0.010 (0.87) (RT Medium), 0.017 (0.97) (PFM), 0.016 (0.98) (DPD - Rigid) and 0.017 (0.97) (DPD - Flexible). The error bars are the unbiased standard deviation of the mean for several randomly generated matrices.

As shown in Fig. 11 the cell velocity increases linearly with the pore size for both experimental and computational models. For small pore sizes the cells start to have MMP-dependent migration, degrading the fibres and opening space for migration. In our model the cell becomes trapped for higher pore sizes than in the experimental setup due to the lack of this mechanism (Fig. 11). For the DPD model with flexible fibres (Fig. 11 **b)**) we observe a higher velocity than in the rigid case, which is expected since matrix elasticity favors migration.

In the experimental assay the authors had also repeated the migration study isolating the role of MMPs. For this purpose, the cells were treated with the potent broad spectrum MMP inhibitor GM6001, also known as ilomastat, which mitigates the influence of MMP activity during migration. In Fig. 12 we present the migration velocity as a function of the pore cross section for **(a)** PFM and **(b)** DPD results and compare with the GM6001 experimental data. As before, we can observe for the three cases a linear dependency with the pore cross section.

**Figure 12.**
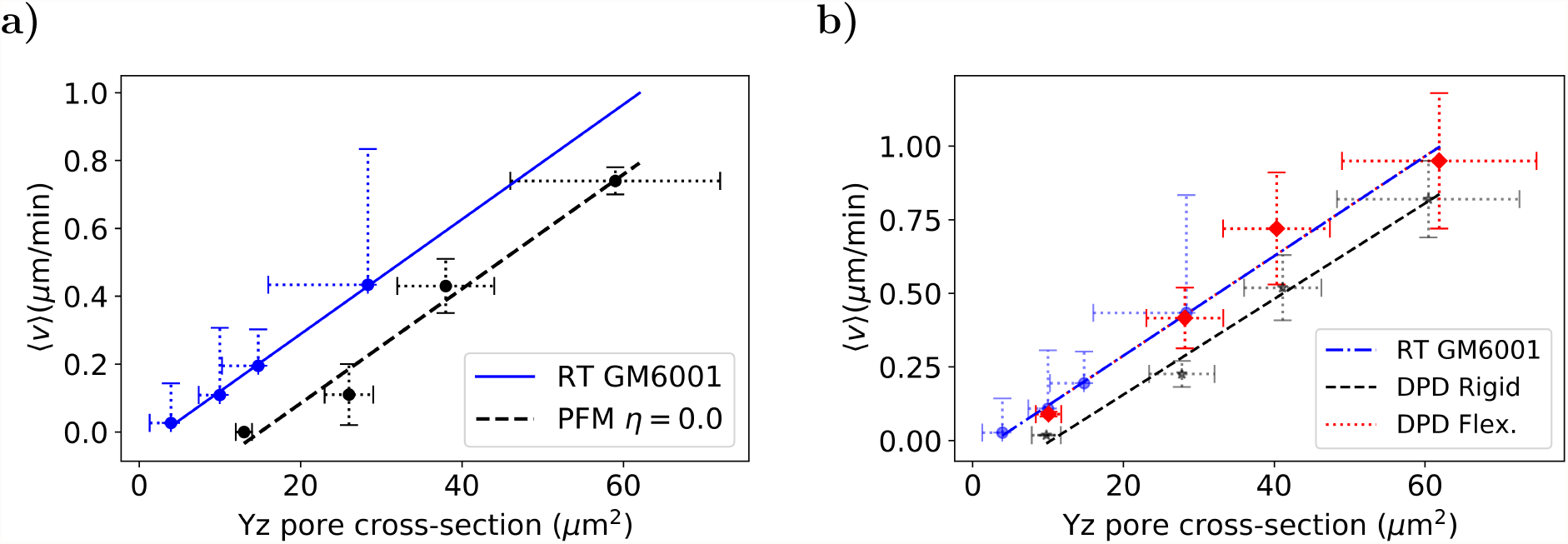
Migration velocity as a function of the pore cross-section *in silico* and *in vitro* [9]. The DPD flexible fitting is perfectly superposed with the RT GM6001 experimental line. The slopes and *R*^2^ are the following: 0.017 (0.99) (RT GM6001), 0.017 (0.97) (PFM), 0.016 (0.98) (DPD - Rigid) and 0.017 (0.97) (DPD - Flexible). The error bars are the unbiased standard deviation of the mean for several randomly generated matrices.

Comparing the two lines in Fig. 12 **a**) we can notice that one is shifted in relation to another, as well when comparing the DPD Rigid in **b**). This difference is the result of a larger plasticity of the cancer cells used in the experiments and their capability of adhering to the fibres, which facilitates migration, when compared with the capacity of the simulated cells to deform. The fibrosarcoma cells are able to migrate under extreme confinement conditions as seen in the transwell filter assay where cells can pass through apertures of only 3 *µ*m wide (Fig. 7-B of [9]). The fact that the line presented in Fig. 12 **b**) for the DPD flexible is shifted to the left corroborate this hypothesis and reveals some level of equivalence between matrix elasticity and cell plasticity.

We then increased the adhesiveness in the PFM and ran simulations for *η* = 0.5 and *η* = 1.0 in order to evaluate the contribution of adhesion to migration. We can see in Fig. 13 that increasing adhesion from *η* = 0 to *η* = 0.5 the velocity dependency with the pore size rotates, becomes more similar to the result obtained experimentally with MMP inhibition (RT GM6001). When increasing even further the adhesiveness (*η* = 1.0) the PFM results became analogous to the experimental results without the inhibitor (RT Medium). These results indicate that MMP inhibitors are able to affect significantly adhesiveness by suppressing the cleavage of molecules that act as adhesion promoters. Moreover this brings adhesiveness as a key modulator of migration in crowded media. In addition, we suggest that more experimental assays should be carried to unveil how MMPs activity acts in different mechanisms of cell migration, using specific inhibitors to target isolated members of the MMP family. Some MMP isoforms are known to affect mainly collagen degradation and remodelling while others act regulation cell adhesion and migration.

**Figure 13.**
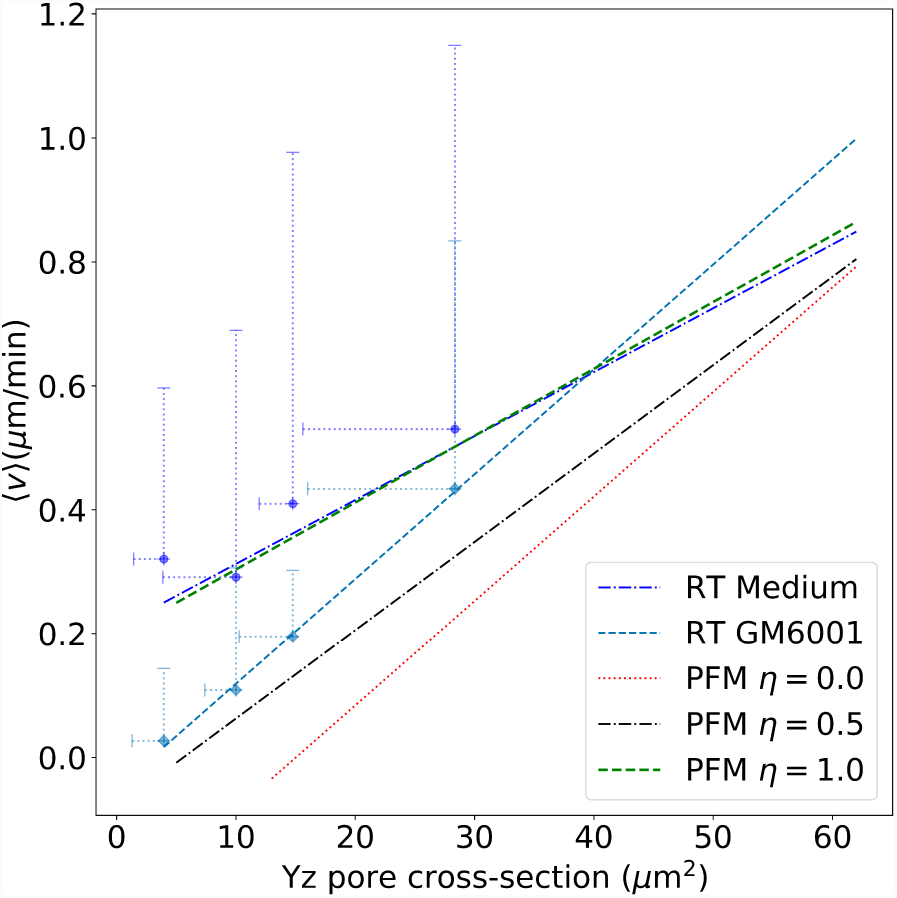
Migration velocity as a function of the pore cross-section for three values of adhesion coefficient *in silico η* = {0, 0.5, 1} and *in vitro* fibrosarcoma cells in rat-tail collagen [9]. The slopes and *R*^2^ for the simulation results from lower to higher adhesions are 0.017 (0.97), 0.014 (0.92), 0.011 (0.92) and 0.010 (0.87) (RT Medium) and 0.017 (0.99) (RT GM6001) for the experimental data.

## 4. Conclusion

In this article we presented two minimalist mathematical models for droplets: the DPD and the PFM. We carried out a systematic study for the movement of a droplet in fibrous media, measuring its migration velocity and morphology while varying the fibre density and adhesiveness. We observed that cell migration is strongly modulated by spatial conditions, and specially by the adhesion between the cell and the substrate.

Our results indicate that adhesion is critical for cell migration [92], by modulating cell morphology [93, 94, 95, 96] in crowded environments and enhancing cell velocity [97]. We have compared our results to experimental data [9, 86]. The theoretical results for the velocity and the experimental data shown the same qualitative behaviour as a function of the matrices’ pore cross section and adhesion strength. The results are model independent and have shown good agreement between the two methodologies and experiments in the literature, which indicates that these minimalist descriptions are able to capture the main features and the basic mechanical nature of cell migration. In addition, the two approaches are complementary to each other, which helps to fulfil some gaps due to model assumptions.

Our analysis suggests that the MMP family plays an important role as adhesion regulator, since our results reproduce the effect of MMP inhibition by the only mean of varying the cell-ECM adhesion: a stronger adhesion in our model gives similar results to the migration assay without MMP inhibition; while a weaker adhesion describes the experimental data with the GM6001 inhibitor. In fact, in the recent years the MMPs have been gathering attention by its many faceted action on modulating cell migration and invasiveness, pushing its importance beyond a collagen degrading enzyme [45, 98, 99]. For example, the MT1-MMP binds more than one hundred of cell surface partners [100, 101], including: tetraspanins family [102], known for regulating cancer progression [100]; integrins and CD44 [103], which are protagonists in cell adhesion [104].

The DPD model is a promising novel approach to model cells in a flexible substrate. In this work we described the fibres as polymeric chains under a simple harmonic potential, but the DPD implementation provides a natural method to introduce nonlinear elasticity. The elastic fibres model allows to explore different problems in cellular systems, such as how cell-to-cell mechanical communication happens through force propagation via the network and its influence on migration. For these reasons further investigation will be carried in order to address the effect of cooperative migration in the tumor microenvironment of metastasis (TMEM).

These results help to elucidate the behaviour of soft cells diffusing in complex media. Mainly, it has shown that adhesion is a key modulator for cellular plasticity and migration in confined conditions. In addition we showed that there is a strong correlation between migration velocity and the cell deformation. We propose that new assays should be carried to address the role of adhesion and the effect of different MMPs in cell migration under confined conditions.

## Acknowledgments

The simulations were done in the Satolep Cluster of the Federal University of Pelotas and in the Centaurus cluster of the University of Coimbra. This work was funded by FEDER funds through the Operational Programme Competitiveness Factors - COMPETE and by national funds by FCT - Foundation for Science and Technology under the strategic projects UID/FIS/04564/2016 and POCI-01-0145-FEDER-031743 – PTDC/BIA-CEL/31743/2017 (MMS, RT). MMS acknowledges the support of the National Council of Technological and Scientific Development (CNPq) under the grant 235101/2014-1 and thanks Pedro V.P. Cunha for the fruitful discussions. The authors would like to thank Beatriz Costa-Gomes for constructive criticism of the manuscript. JRB acknowledges CNPq and FAPERGS for financial support.

## References

[1] Lambert A W, Pattabiraman D R and Weinberg R A 2017 Cell 168 670–691

[2] Taketo M M 2011 Cancer prevention research 4 324–328

[3] Steeg P S 2016 Nature reviews cancer 16 201

[4] Mehlen P and Puisieux A 2006 Nature reviews cancer 6 449

[5] Fidler I J 2003 Nature reviews cancer 3 453

[6] Venning F A, Wullkopf L and Erler J T 2015 Frontiers in oncology 5 224

[7] Lamouille S, Xu J and Derynck R 2014 Nature reviews Molecular cell biology 15 178

[8] Chiodoni C, Colombo M P and Sangaletti S 2010 Cancer and Metastasis Reviews 29 295–307

[9] Wolf K, Te Lindert M, Krause M, Alexander S, Te Riet J, Willis A L, Hoffman R M, Figdor C G, Weiss S J and Friedl P 2013 The Journal of cell biology 201 1069–84 ISSN 1540-8140

[10] Brodland G W and Veldhuis J H 2012 PLoS One 7 e44281

[11] Malandrino A, Kamm R D and Moeendarbary E 2017 ACS biomaterials science & engineering 4 294–301

[12] Katira P, Bonnecaze R T and Zaman M H 2013 Frontiers in oncology 3 145

[13] Wise S M, Lowengrub J S, Frieboes H B and Cristini V 2008 Journal of theoretical biology 253 524–543

[14] Byrne H and Chaplain M A 1996 Mathematical and Computer Modelling 24 1–17

[15] Hay E D 2013 Cell biology of extracellular matrix (Springer Science & Business Media)

[16] Frantz C, Stewart K M and Weaver V M 2010 J Cell Sci 123 4195–4200

[17] Friedl P and Alexander S 2011 Cell 147 992–1009

[18] Visse R and Nagase H 2003 Circulation research 92 827–839

[19] Kessenbrock K, Plaks V and Werb Z 2010 Cell 141 52–67

[20] Wolf K and Friedl P 2011 Trends in cell biology 21 736–744

[21] Khatiwala C B, Peyton S R and Putnam A J 2006 American Journal of Physiology-Cell Physiology 290 C1640–C1650

[22] Goetz J G, Minguet S, Navarro-Lérida I, Lazcano J J, Samaniego R, Calvo E, Tello M, Osteso-Ibáñez T, Pellinen T, Echarri A et al. 2011 Cell 146 148–163

[23] Wullkopf L, West A K V, Leijnse N, Cox T R, Madsen C D, Oddershede L B and Erler J T 2018 Molecular biology of the cell mbc–E18

[24] Gaggioli C, Hooper S, Hidalgo-Carcedo C, Grosse R, Marshall J F, Harrington K and Sahai E 2007 Nature cell biology 9 1392

[25] Lugassy C, Zadran S, Bentolila L A, Wadehra M, Prakash R, Carmichael S T, Kleinman H K, Péault B, Larue L and Barnhill R L 2014 Cancer Microenvironment 7 139–152

[26] Wang W, Wyckoff J B, Goswami S, Wang Y, Sidani M, Segall J E and Condeelis J S 2007 Cancer research 67 3505–3511

[27] Sahai E, Wyckoff J, Philippar U, Segall J E, Gertler F and Condeelis J 2005 BMC biotechnology 5 14

[28] Yamauchi K, Yang M, Jiang P, Yamamoto N, Xu M, Amoh Y, Tsuji K, Bouvet M, Tsuchiya H, Tomita K et al. 2005 Cancer research 65 4246–4252

[29] Weigelin B, Bakker G J and Friedl P 2012 IntraVital 1 32–43

[30] Provenzano P P, Eliceiri K W, Campbell J M, Inman D R, White J G and Keely P J 2006 BMC medicine 4 38

[31] Paul C D, Mistriotis P and Konstantopoulos K 2017 Nature Reviews Cancer 17 131

[32] Friedl P and Wolf K 2010 The Journal of cell biology 188 11–19

[33] Friedl P 2004 Current opinion in cell biology 16 14–23

[34] Costa P and Parsons M 2010 New insights into the dynamics of cell adhesions International review of cell and molecular biology vol 283 (Elsevier) pp 57–91

[35] Alberts B, Bray D, Lewis J, Raff M, Roberts K and Watson J 2002 Inc., London

[36] Zaidel-Bar R, Cohen M, Addadi L and Geiger B 2004 Hierarchical assembly of cell–matrix adhesion complexes

[37] Destaing O, Saltel F, Géminard J C, Jurdic P and Bard F 2003 Molecular biology of the cell 14 407–416

[38] Oikawa T and Takenawa T 2009 Cell adhesion & migration 3 195–197

[39] Jabłońska-Trypuć A, Matejczyk M and Rosochacki S 2016 Journal of enzyme inhibition and medicinal chemistry 31 177–183

[40] Gross J and Lapiere C M 1962 Proceedings of the National Academy of Sciences of the United States of America 48 1014

[41] Rodríguez D, Morrison C J and Overall C M 2010 Biochimica et Biophysica Acta (BBA)-Molecular Cell Research 1803 39–54

[42] Jiao Y, Feng X, Zhan Y, Wang R, Zheng S, Liu W and Zeng X 2012 PloS one 7 e41591

[43] Conant K, T Lim S, Randall B and A Maguire-Zeiss K 2012 Current HIV research 10 384–391

[44] Butler G S, Dean R A, Tam E M and Overall C M 2008 Molecular and cellular biology 28 4896–4914

[45] Ferrari R, Martin G, Tagit O, Guichard A, Cambi A, Voituriez R, Vassilopoulos S and Chavrier P 2019 Nature communications 10 1–15

[46] Ubezio B, Blanco R A, Geudens I, Stanchi F, Mathivet T, Jones M L, Ragab A, Bentley K and Gerhardt H 2016 Elife 5 e12167

[47] Li X, Padhan N, Sjöström E O, Roche F P, Testini C, Honkura N, Sáinz-Jaspeado M, Gordon E, Bentley K, Philippides A et al. 2016 Nature communications 7 11017

[48] Osborne J M, Fletcher A G, Pitt-Francis J M, Maini P K and Gavaghan D J 2017 PLoS computational biology 13 e1005387

[49] Lakatos D, Somfai E, Méhes E and Czirók A 2018 Journal of theoretical biology 456 261–278

[50] Frieboes H B, Jin F, Chuang Y L, Wise S M, Lowengrub J S and Cristini V 2010 Journal of theoretical biology 264 1254–1278

[51] Lima E, Oden J and Almeida R 2014 Mathematical Models and Methods in Applied Sciences 24 2569–2599

[52] Lorenzo G, Hughes T J, Dominguez-Frojan P, Reali A and Gomez H 2019 Proceedings of the National Academy of Sciences 116 1152–1161

[53] Travasso R D, Castro M and Oliveira J C 2011 Philosophical Magazine 91 183–206

[54] Milde F, Bergdorf M and Koumoutsakos P 2008 Biophysical journal 95 3146–3160

[55] Santos-Oliveira P, Correia A, Rodrigues T, Ribeiro-Rodrigues T M, Matafome P, Rodríguez-Manzaneque J C, Seiça R, Girão H and Travasso R D M 2015 PLoS Comput Biol 11 e1004436

[56] Moreira-Soares M, Coimbra R, Rebelo L, Carvalho J and Travasso R D M 2018 Scientific Reports 8 1–12

[57] Vilanova G, Colominas I and Gomez H 2017 Journal of The Royal Society Interface 14 20160918

[58] Palmieri B, Bresler Y, Wirtz D and Grant M 2015 Scientific Reports 5 11745

[59] Najem S and Grant M 2016 Physical Review E 93 052405

[60] Najem S and Grant M 2015 Soft matter 11 4476–4480

[61] Najem S and Grant M 2014 Soft matter 10 9715–9720

[62] Najem S and Grant M 2013 Physical Review E 88 034702

[63] Shao D, Levine H and Rappel W j 2012 Proceedings of the National Academy of Sciences of the United States of America 109 6851–6 ISSN 1091-6490

[64] Marth W and Voigt A 2014 Journal of Mathematical Biology 69 91–112 ISSN 14321416

[65] Marth W, Praetorius S and Voigt A 2015 Journal of The Royal Society Interface 12 20150161

[66] Kulawiak D A, Camley B A and Rappel W J 2016 Plos Comp Biol ISSN 1553-7358

[67] Camley B A, Zhao Y, Li B, Levine H and Rappel W J 2017 Physical Review E 95 012401

[68] Moure A and Gomez H 2017 Computer Methods in Applied Mechanics and Engineering 320 162–197

[69] Groot R D and Rabone K L 2001 Biophysical Journal 81 725–736

[70] Yamamoto S, Maruyama Y and Hyodo S a 2002 The Journal of chemical physics 116 5842–5849

[71] Basan M, Prost J, Joanny J F and Elgeti J 2011 Physical biology 8 026014

[72] Ye T, Phan-Thien N and Lim C T 2016 Journal of biomechanics 49 2255–2266

[73] Hoogerbrugge P and Koelman J 1992 EPL (Europhysics Letters) 19 155

[74] Nonomura M 2012 PLoS ONE 7 1–9 ISSN 19326203

[75] Akiyama M, Nonomura M, Tero A and Kobayashi R 2018 Physical biology 16 016005

[76] Bray A 1994 Advances in Physics

[77] Bray A J, Cavagna A and Travasso R D 2001 Physical Review E 65 016104

[78] Moreira-Soares M 2019 Software package for cell research (spiccato) URL https://doi.org/10.5281/zenodo.3534284

[79] Moreira-Soares M 2019 Trajpy URL https://doi.org/10.5281/zenodo.3537983

[80] Liu M, Liu G, Zhou L and Chang J 2015 Archives of Computational Methods in Engineering 22 529–556

[81] Macis M, Lugli F and Zerbetto F 2017 ACS applied materials & interfaces 9 19552–19561

[82] Allen M P and Schmid F 2007 Molecular Simulation 33 21–26

[83] Lobaskin V and Dünweb B 2004 New Journal of Physics 6 54

[84] Blumers A L, Tang Y H, Li Z, Li X and Karniadakis G E 2017 Computer Physics Communications 217 171 – 179 ISSN 0010-4655

[85] Gatsonis N A, Potami R and Yang J 2014 Journal of Computational Physics 256 441 – 464 ISSN 0021-9991

[86] Dieterich P, Klages R, Preuss R and Schwab A 2008 Proceedings of the National Academy of Sciences of the United States of America 105 459–63 ISSN 1091-6490

[87] Maheshwari G, Wells A, Griffith L G and Lauffenburger D A 1999 Biophysical Journal 76 2814–2823 ISSN 00063495

[88] Kelley L C, Lohmer L L, Hagedorn E J and Sherwood D R 2014 Traversing the basement membrane in vivo: A diversity of strategies

[89] Maheshwari G, Brown G, Lauffenburger D A, Wells A and Griffith L G 2000 Journal of cell science 113 (Pt 1 1677–86 ISSN 0021-9533

[90] Bordin J R 2018 Physica A: Statistical Mechanics and its Applications 495 215–224

[91] Bordin J R 2019 Fluid Phase Equilibria 499 112251

[92] Lock J G, Wehrle-Haller B and Strömblad S 2008 Cell–matrix adhesion complexes: master control machinery of cell migration Seminars in cancer biology vol 18 (Elsevier) pp 65–76

[93] Natale C, Lafaurie-Janvore J, Ventre M, Babataheri A and Barakat A 2019 Journal of the Royal Society Interface 16 20190263

[94] Oakes P W, Banerjee S, Marchetti M C and Gardel M L 2014 Biophysical journal 107 825–833

[95] Bischofs I B, Schmidt S S and Schwarz U S 2009 Physical review letters 103 048101

[96] Banerjee S and Marchetti M C 2013 New Journal of Physics 15 035015

[97] Tantivejkul K, Vucenik I and Shamsuddin A M 2003 In vivo 10 12

[98] Takino T, Watanabe Y, Matsui M, Miyamori H, Kudo T, Seiki M and Sato H 2006 Experimental cell research 312 1381–1389

[99] Itoh Y 2006 IUBMB life 58 589–596

[100] Tomari T, Koshikawa N, Uematsu T, Shinkawa T, Hoshino D, Egawa N, Isobe T and Seiki M 2009 Cancer science 100 1284–1290

[101] Knapinska A M and Fields G B 2019 Pharmaceuticals 12 ISSN 1424-8247

[102] Schröder H, Hoffmann S, Hecker M, Korff T and Ludwig T 2013 The international journal of biochemistry & cell biology 45 1133–1144

[103] Gálvez B G, Matías-Román S, Yáñez-Mó M, Sánchez-Madrid F and Arroyo A G 2002 The Journal of cell biology 159 509–521

[104] Senbanjo L T and Chellaiah M A 2017 Frontiers in cell and developmental biology 5 18

