## Supporting Information. for "Adhesion modulates cell morphology and migration within dense fibrous networks"

### Supporting Information for:
### *"Adhesion modulates cell morphology and migration within dense fibrous networks"*


Maurício Moreira-Soares,[1,*] Susana P. Cunha,[2]
José Rafael Bordin,[3,†] and Rui D. M. Travasso[1]

[1] *CFisUC, Department of Physics, University of Coimbra, Rua Larga, 3004-516 Coimbra, Portugal*

[2] *CQC, Department of Chemistry, University of Coimbra, Rua Larga, 3004-535 Coimbra, Portugal*

[3] *Department of Physics, Institute of Physics and Mathematics, Federal University of Pelotas, Rua dos Ipês, Capão do Leão, RS, 96050-500, Brazil*



(Dated: November 11, 2019)



*

†

#### S1. TRAJECTORY PROFILE IN THE PHASE-FIELD MODEL

In order to analyse the cell movement in the PFM we tracked the position of the cell's center of mass along the time. In the Fig. 1 we have three trajectories for $\rho = 0.20,\ 0.55,\ 0.60$ and for each value of adhesion between the cell and the fibres $\eta = 0,\ 1$. We can observe that for $\eta = 0$ and $\rho = 0.60$ the cell has longer freezing periods due to the difficulty of migrating through smaller pores. When we increase adhesion the cell movement is facilitated and the moments the cell is stuck are almost imperceptive. Moreover, the profiles presented for the phase-field model are similar to the ones shown in Fig. 6 of the main text for the DPD model.

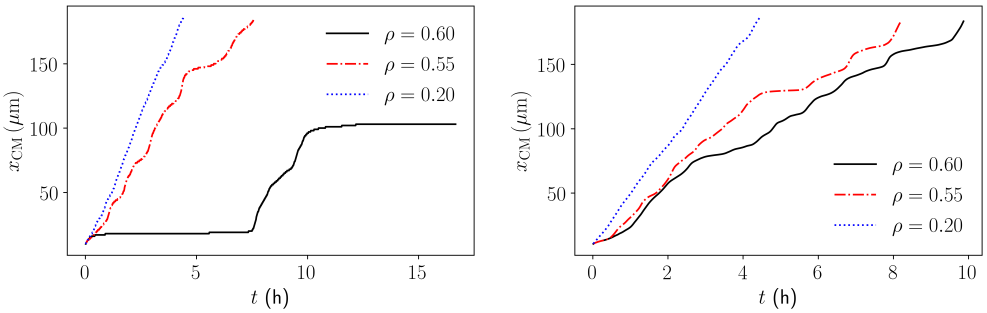


**a)** $\eta = 0$ **b)** $\eta = 1.0$

FIG. 1. Typical cell trajectories in the PFM for two values of adhesion $\eta = \{0, 1\}$ and three densities $\rho = \{0.20, 0.55, 0.60\}$ taking into consideration only the position of the center of mass in the $x-$direction. For higher adhesions the cell presents longer trapped times. However, when we increase the adhesion it is noticeable the improvement in migration, in special for $\rho = 0.60$ where the trapped periods decrease considerably.

#### S2. MATRIX STIFFNESS

We have measured the velocity as function of the repulsion coefficient $\gamma$ for the PFM and as function of the fibres' rigidity $\kappa$ for the DPD model. The coefficient $\gamma$ is related to how strong the depletion interaction acts to avoid an overlap between the cell and the fibres, and it is akin to the effect by matrix stiffness. For the DPD model the stiffness is introduced in the description of the fibres as polymeric chains of monomers that interact through a harmonic potential with spring constant $\kappa$. Fig. 2 shows a similar velocity dependency as a function of $\gamma^{-1}$ and $\kappa^{-1}$: the velocity decreases for higher values of $\gamma$ and $\kappa$, and it increases as we decrease these coefficients. Simulations were performed with $\kappa$ varying from 10 N/$\mu$m to $\infty$ in order to check the dependency of the cell velocity with the chain flexibility.

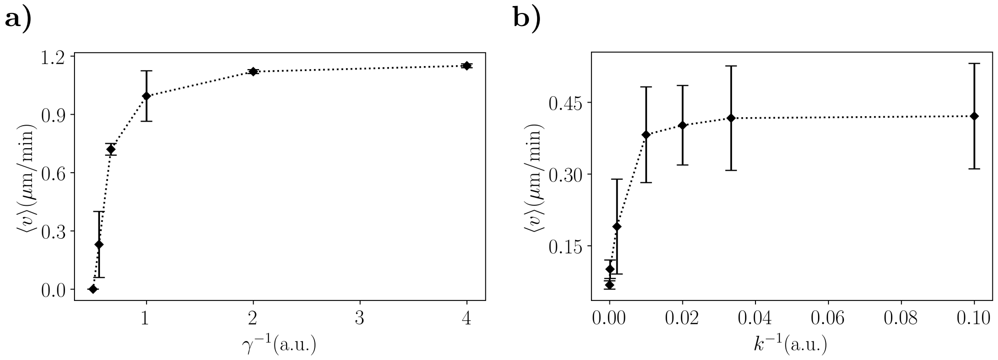


FIG. 2. Cell velocity in the matrix as function of a) the inverse of the repulsion coefficient $\gamma^{-1}$ and b) of the inverse of the fibre rigidity $\kappa^{-1}$ for density $\rho = 0.70$ and adhesiveness $\eta = 0$.

#### S3. COMPUTATIONAL METHODS

##### Simulations

The simulations for the PFM were performed in the Navigator Cluster of the University of Coimbra. The simulation box had the dimensions $250 \times 37.5 \times 37.5\,\mu$m and periodic boundary conditions were applied. We have implemented the finite differences method with central differences for the computation of the Laplace operator and Euler's scheme for the time evaluation. The main code was written in Fortran 2003 and the python scripts for data analysis are available in the online repository https://phydev.github.io/SPiCCAto [1]. We used the package TrajPy for trajectory classification [2]. For each set of parameters we ran 10 simulations with randomly distributed fibres using the pseudorandom number generator *ran2*.

In the DPD model the simulation box is a parallelepiped with size 25.0 $\mu$m $\times$ 25.0 $\mu$m $\times$ 125.0 $\mu$m in the $x$, $y$ and $z$ directions, respectively. Periodic boundary conditions are applied in all directions. The temperature was fixed at 300 K, and the relation $T = 133T^{*}+240$ [3, 4] was used to obtain the reduced temperature. The unit of length was set to $\sigma = 1.25\mu$m, the damping parameter was $\gamma = 5.0$ and the time step used was $0.02\tau$, where $\tau$ is the time scale.

The fibres occupy a region of 100.0 $\mu$m in the $z$-direction, with two 12.5 $\mu$m buffers at the beginning and at the end of the simulation box in this direction. Initially, the cell was placed in the left buffer and we equilibrated the system for $5 \times 10^6$ steps. During the equilibration time the ghost bead is fixed to ensure that the cell does not diffuse. Afterwards, a force was applied to the cell to mimic the flow along the chemotactic gradient. This force is constant, with intensity $1 \times 10^{-11}$ N along the $z$-direction. Then we analysed the system properties during the migration in the fibre network region. When the cell reaches the right buffer the simulation ends. This was repeated 10 times for each simulated point. We show in Fig. 4 of the main document a schematic depiction of the simulation box.

To obtain the time scale to evaluate the velocity we employed the procedure proposed recently by Macis and co-workers [3]. We evaluate the self-diffusion coefficient of the cell in equilibrium bulk simulations. The value obtained was $D^* = 0.01566\sigma^2/\tau$. Then, we can estimate the time scale using the relation [3],

$$\tau = \frac{D^*}{D}\sigma^2 = 15.66s \tag{1}$$

where we have used $D \approx 10^{-15}\text{m}^2/\text{s}$ as the experimental self-diffusion coefficient of an endothelial cell [5].

**Pore Cross-Section estimation**

In this work we used two different clustering algorithms to determine the pore cross-section of our matrices. For the matrices in the PFM we applied the Connected Component Algorithm (CCA) implemented in the machine learning library Scikit-Learn (See Fig. 3) [6]. The CCA verifies the connectivity between the pixels and classifies each contiguous region surrounded with material as a pore, assigning a unique identifier to each pore. At the end we can easily estimate the area of the pores.

Each matrix was sliced in several planes that are orthogonal to the migration direction and for each plane the CCA was applied to obtain the area of the pores. Then the procedure was repeated for several random generated matrices with the same density and the mean values and their respective unbiased standard deviation were obtained.

On the other hand, for the DPD matrices we used the algorithm presented in the Ref. [7] where the effective area is used to estimate the available empty space. To characterise the

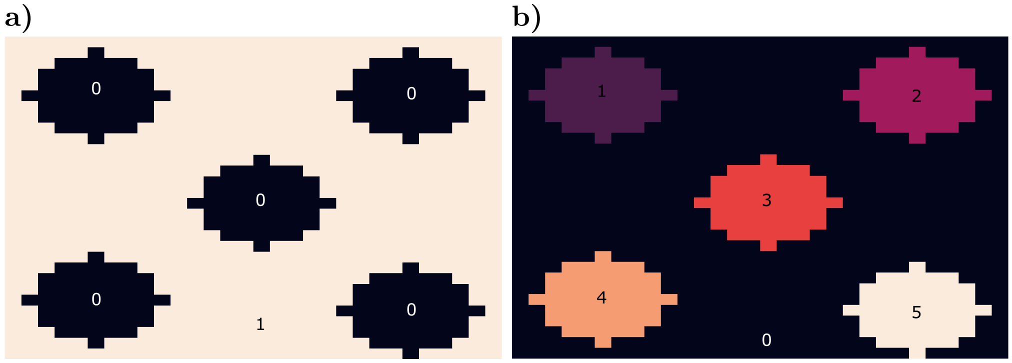


FIG. 3. The clustering algorithm identifies each connected region and then assigns the respective label to each point. a) The initial image contains the pores as zeros and the matrix as ones. b) After applying the CCA each pore is labeled individually with a unique number and the substrate is labeled with zeros.

pores we took a collection of 1000 snapshots of each simulation run and attempted to insert ghost particles with diameter 1.0 $\mu$m in the fibre matrix lattice (the center of those ghost particles were allowed to be in a lattice with side 0.25$\mu$m). If there are no overlaps with the polymers or with ghost particles already inserted, a new ghost particle is inserted in the chosen position. We consider a pore when the hole is filled with at least 2 ghost particles. Pores smaller than this were considered as small defects and were not taken into account for the analysis. With this, we can use the number particles in each ghost particle cluster, as a proxy of the pore size $A_p$.

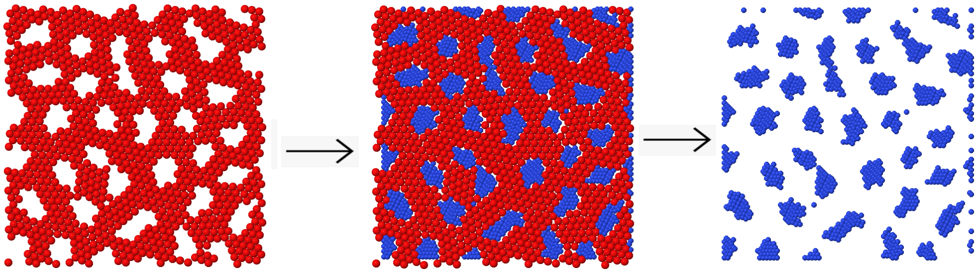

FIG. 4. Schematic depiction of the pore area evaluation. First, we select a snapshot of the system. Here, the fibres are the red spheres, and a 2D projection is shown for simplicity. Then, we attempt to insert ghost particles. If there is no overlap, a ghost (blue sphere) is inserted. With the positions of the ghost particles we can evaluate the pore's properties.

#### S4. PARAMETERS

##### Phase-Field Model

The parameters used in the model are presented in the Table I. The values were chosen based on biological measures available in the literature.

TABLE I. **Description of the parameters used in the PFM.**

| Parameter | Description | Value *in Silico* | *in vivo* |
|---|---|---|---|
| $a$ | Lattice parameter | 1 | $1.25\,\mu$m |
| $\Delta t$ | Unity of time | 1 | 1 min |
| $R_d$ | Cell diameter | 12 | $15\,\mu$m |
| $D_f$ | Fibre diameter | 2 | $2.5\,\mu$m |
| $\varepsilon$ | Interface cost | 1 | – |
| $\delta$ | Interface width | $\mathcal{O}(4\varepsilon)$ | – |
| $\sigma$ | Surface tension | $\varepsilon/6\sqrt{2}$ | – |
| $\eta$ | Adhesion strength | $0.0 \sim 1.5$ | – |
| $\gamma$ | Repulsion strength | $1.0 \sim 2.0$ | – |
| $\chi$ | Chemotactic coefficient | 0.80 | $1.2\,\mu$m/min |
| $\rho$ | Density of fibres | $0 \sim 1(V_{\text{fibres}}/V_{\text{total}})$ | – |

##### Dissipative Particle Dynamics

The parameters used in the DPD model were chosen with the aim to assure that the adhesion energy and the interface velocity would have similar profiles as in the PFM. The Table II presents the list of parameters used in the DPD model.

TABLE II. **Description of the parameters used in the DPD model.**

| **Parameter** | **Description** | **Value *in Silico*** | ***in vivo*** |
|---|---|---|---|
| $\tau$ | Time scale | $D^*/D\sigma^2$ | 15.66 s |
| $\delta t$ | Time step | $0.02\tau$ | 0.31 s |
| $t_{eq}$ | Equilibration time | $5 \times 10^6$ | – |
| $T$ | Temperature | – | 300K |
| $a_c$ | Cell radius | – | 6 $\mu$m |
| $a_{cc}$ | Cell beads depletion strength | – | $12.5 \times 10^{-11}$ N |
| $\rho_N$ | Solvent number density | 4 | – |
| $a_{cs}$ | Cell-Solvent repelling strength | – | $65 \times 10^{-11}$ N |
| $b_{cf}$ | Cell-fibre adhesion coefficient | $\eta_{DPD} a_{cf}$ | – |
| $k_{cg}$ | Cell-ghost bead coupling | – | 30 N/m |
| $k_{ff}$ | Fibre rigidity | – | 30 N/m |
| $r_1$ | Cut-off distance for repulsion | – | 1.25$\mu$m |
| $r_2$ | Cut-off distance for attraction | – | 2.50$\mu$m |
| $\gamma$ | Dampening | 5.0 | – |
| $D$ | Diffusion constant | – | $10^{-15}$ m$^2$/s |
| $F_\chi$ | Chemotactic force | – | $1 \times 10^{-11}$ N |

#### REFERENCES


[1] M. Moreira-Soares, “Software package for cell research (spiccato),” (2019).

[2] M. Moreira-Soares, “Trajpy,” (2019).

[3] M. Macis, F. Lugli, and F. Zerbetto, ACS applied materials & interfaces **9**, 19552 (2017).

[4] X. Li, I. V. Pivkin, H. Liang, and G. E. Karniadakis, Macromolecules **42**, 3195 (2009).

[5] J. A. Sherratt and J. D. Murray, Proceedings of the Royal Society of London B: Biological Sciences **241**, 29 (1990).

[6] F. Pedregosa, G. Varoquaux, A. Gramfort, V. Michel, B. Thirion, O. Grisel, M. Blondel, P. Prettenhofer, R. Weiss, V. Dubourg, J. Vanderplas, A. Passos, D. Cournapeau, M. Brucher, M. Perrot, and E. Duchesnay, Journal of Machine Learning Research **12**, 2825 (2011).
[7] J. R. Bordin, Physica A: Statistical Mechanics and its Applications **495**, 215 (2018).
